# Modelling the On-going Natural Selection of Educational Attainment in Contemporary Societies

**DOI:** 10.1101/605311

**Authors:** Ze Hong

**Affiliations:** Department of Human Evolutionary Biology, Harvard University, 11 Divinity Avenue, 02138 Cambridge MA, United States

## Abstract

There has been substantial increase in education attainment (EA) in both developing and developed countries over the past century. I present a simulation model to examine the potential evolutionary trajectories of EA under current selective pressure in western populations. With the assumption that EA is negatively correlated with fitness and has both a genetic component and a cultural component, I show that when prestige-biased transmission of the EA (i.e. people with more education are more likely to be copied) is present, the phenotype of EA is likely to keep increasing in the short term, yet the genetic component of EA may be undergoing a constant decline and eventually lead to an overall decrease in the phenotype.

**Significance statement:** Contemporary humans live in very different environments from our ancestors and are subject to very different selective forces. The future evolutionary trajectory of current human populations is thus of both intellectual and pragmatic importance. The present simulation model sheds light on the short-term evolutionary dynamics of educational attainment, and demonstrates the significant contribution of systematically transmitted human culture in shaping the evolutionary paths and producing non-intuitive evolutionary outcomes.

## Introduction

Today, the importance of taking evolutionary perspectives in understanding physical and physiological traits of humans is well-recognized (1–3), and there is little doubt that humans have experienced selection in relatively recent human evolutionary history (4). Indeed, many human physical and physiological trait variations have been celebrated as evidence of natural selection or gene-culture coevolution (5). Increasingly, researchers in the social sciences are paying more attention to evolutionary explanations for universality in human behavioral and cognitive traits (6, 7).

Nonetheless, to this day the majority of evolutionary studies on *homo sapiens* have focused on the past, using evolutionary theories to explain evolved physical, cognitive, and behavioral traits (8, 9), with the notable exception of human behavioral ecology, which primarily focuses on human phenotypic plasticity in small scale societies within the fitness-maximization framework (10, 11). However, recent advances in genomic sciences and statistical techniques provide us with a better understanding of the genetic architecture of quantitative traits in humans, which, combined with observational studies on how selective pressure is operating in contemporary human populations, makes it possible to detect on-going selection and to potentially predict the future evolutionary trajectory of specific traits (12–14). Of course, special attention needs to be paid when behavioral traits are under consideration, as the transmission of human culture could systematically and significantly affect the evolutionary dynamics (15).

Educational attainment (EA) has long interested social scientists. It is strongly, negatively associated with fertility (16, 17) and extensive research in behavioral genetics estimates significant heritability (18, 19). Large scale Genome Wide Association Studies have identified specific loci that contribute to EA (20, 21), and two recent studies show that in certain western populations there is on-going negative selection against EA (13, 22), where the genetic component of EA as measured by polygenic score has been declining for the past century in Iceland and the US respectively. On the other hand, the phenotype of EA, along with other cognitive measures such as the IQ score, has drastically increased in western populations (23). To explore the unusual evolutionary dynamics of EA, I construct an agent based simulation model to track the relative change in both genotypic value and phenotypic value of EA assuming that 1) EA is negatively associated with fitness; 2) the phenotype of EA has both a genetic and a cultural component; 3) the genetic component is vertically transmitted from parents to offspring and the cultural component is obliquely transmitted (24) from the parental to the offspring generation and 4) the cultural component is transmitted with a prestige-based bias, which refers to the psychological tendency of humans to copy the behaviors of the more prestigious individuals in a community (25). Alternatively, the tendency to copy from more educated people can be viewed as payoff biased transmission if education itself is considered as a desirable outcome.

The present simulation model differs from traditional evolutionary models in two ways. First, this model is not aiming at identifying evolutionarily stable strategies (26); it is rather an attempt to combine standard evolutionary logic and cultural evolutionary theories (27) to explain and potentially predict genetic and phenotypic changes of a specific trait (EA) over time. Second, this model explicitly specifies the phenotype (EA) function as having a gene-culture interaction term in addition to the additive components of gene and culture. Orthodox gene-culture coevolution theory explicitly recognizes the importance of both factors, but typically models the phenotypic contribution of gene and culture in an additive fashion (15), presumably for analytical tractability. In a recent proposal, Day and Bonduriansky (2011) provide a theoretical framework to account for both genetic and non-genetic inheritance under a generalized from of the Price equation (29). However, to quantitatively track evolutionary dynamics they also assume that the phenotype is additively determined by a genetic component and a non-genetic component. As will be seen, the inclusion of a gene-culture interaction term significantly alters the evolutionary trajectories of EA.

## Methods

I employ the common assumptions of asexual reproduction and non-overlapping generations in theoretical evolution (15, 28). Each individual is represented by a vector [g, z] where g stands for the genotypic value, and p stands for the phenotypic value (educational attainment). In all simulation runs, the population is initially seeded with individuals drawn from the uniform distribution (0,1). “Cultural fitness”, or the relative likelihood that a particular individual is picked as a model, is defined as z_i_^φ^, where φ is a population level parameter which roughly represents the idea of a “cultural reproductive skew”. When φ>1, individuals with high cultural fitness have disproportionally larger probabilities to produce cultural offspring by serving as models for culturally naïve individuals, and when φ=0, all individuals in the population are equally likely to be picked as a model.

Starting from generation 2, for any given individual i, its phenotype of interest (educational attainment) is defined to be

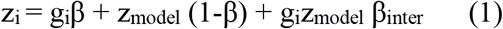

where g_i_β represents the genetic contribution from the focal individual, z_model_ (1−β) represents the cultural contribution from the cultural model that the focal individual picked from the previous generation based on the model’s cultural fitness, and g_i_ z_model_ β_inter_ represents the interaction between the genetic component and the cultural component. I constrain the β coefficients for the first two terms to sum to 1 as it reduces the total number of parameters and I am more interested in the relative contribution from gene and culture and their magnitude compared to β_inter_.

For any given individual i, its genetic fitness w_i_ is defined to be

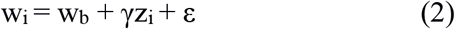

where w_b_ represents the baseline fitness, γz_i_ represents the fitness contribution from the phenotype of interest (EA), and ε is a uniformly distributed error term. Importantly, genetic fitness is forced to be non-negative. In cases where the numeric value of an individual’s genetic fitness becomes negative during simulation runs, it is automatically set to be 0.

In the beginning of the life cycle, an individual is born with a genotypic value inherited from his parent, and will sample the whole parent generation and pick an individual as his model based on cultural fitness. His own phenotypic value is then determined by the phenotype function (1). Once all individuals in the focal generation have acquired a phenotypic value, they become the new parental generation and produce offspring based on their genetic fitness as determined by (2). At the end of each life cycle, both the genotypic value and phenotypic value can change (mutate) with some probability in frequency and magnitude.

To further explore the evolutionary dynamics in relevant contemporary contexts, I added a “demographic shock” condition by having EA uncorrelated with fitness up to a fixed generation number and then changed the correlation into negative. This is to mimic the process of demographic transition which happened in most post-industrialized societies (30, 31) in a relatively short amount of time.

## Results

### General result

The main result from my simulation model is that phenotypic value of EA can increase as the genotypic value of EA decreases (Figure 1, top left plot), at least for a short period of time (~23 generations for benchmark parameter settings). There are two opposing forces at work here: the negative correlation between phenotype (EA) and genetic fitness which constantly drives down genotypic value, and the prestige-biased transmission which increases population mean phenotypic value. After running a large number of simulations I identified two key parameters that contribute to this pattern of initial divergence of genotypic value and phenotypic value. First, increase in phenotype depends on the presence of variation in phenotype. In the model, the main source of phenotypic variation is supplied by random “cultural mutation” at the end of each life cycle. This result is largely intuitive: if every individual in the population has roughly the same phenotypic value, then there is less potential room for biased transmission to increase population mean. Second, the interaction term contributes to mean phenotypic value increase by magnifying the effect of biased transmission, as it places extra weight on the model’s phenotypic value in constructing the focal individual’s phenotype. When β_inter_ is set to be 0, the initial phenotypic increase was much milder (Figure 1, top right plot).

**Fig 1.**
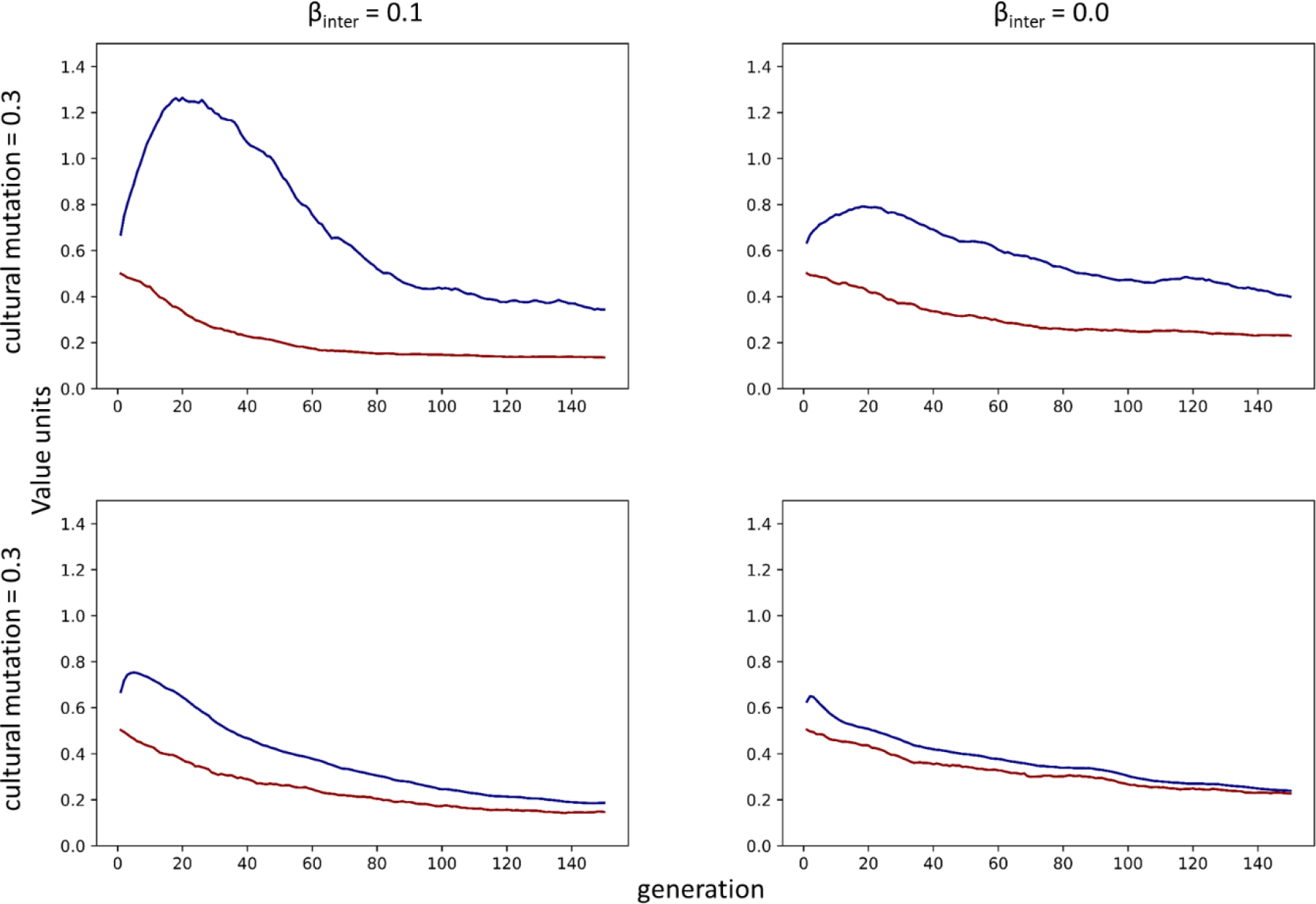
The short term evolutionary trajectory of the genotypic value and phenotypic value of EA. Red lines represent the evolutionary trajectory of genotypic value and blue lines represent the evolutionary trajectory of phenotypic value. The significant diverging pattern (increase in phenotypic value but decrease in genotypic value) occurs only when both cultural mutation and gene-culture interaction term are both present.

As genetic fitness is negatively correlated with phenotype (EA), genotypic value always monotonically decreases from the beginning until variation is exhausted in all simulation runs. The initial increase under certain parameter settings does not last indefinitely and we quickly see a subsequent drop over time because the variance in both genotypic and phenotypic values decreases due to stabilizing selection (32). Eventually, the increase in phenotypic value due to picking a model with higher EA becomes insufficient to compensate for the overall population level decrease in phenotypic value due to natural selection acting against the phenotype.

In the vast majority of simulation runs genotypic value stabilizes after ~ 120 generations. This phenomenon is entirely due to the exhaustion of genetic variance. In the initial population, genotypic values and phenotypic values are randomly seeded, and I assume a very small mutation rate for the genetic component of EA because genetic mutation rate is vastly lower than environmental variations (33). Additionally, the simulation only looks at a relatively short evolutionary time period (~120 generation). As a result, once natural selection has used up all existing genetic variation no further genetic evolution can occur. In cases where variation in environmental contribution (cultural mutation) is large, all observed phenotypic variation would be due to environmental differences (heritability = 0); in cases where there is no variation in environmental contribution, there would be no phenotypic variation nor genotypic variation (i.e. every individual has the same genotypic value and phenotypic value).

### Adding demographic shock

In the “demographic shock” condition, I model educational attainment as first contributing nothing to fitness (γ=0), then after several generations (generation 20 in the figure) EA suddenly becomes negatively associated with fitness (γ=−0.5). As shown in Figure 2, the demographic shock does not immediately drive down average phenotypic value; rather, there is a lag in the phenotype response. Genotypic value, on the other hand, remains constant for the “pre-shock” period and starts to decrease immediately after the shock.

**Fig 2.**
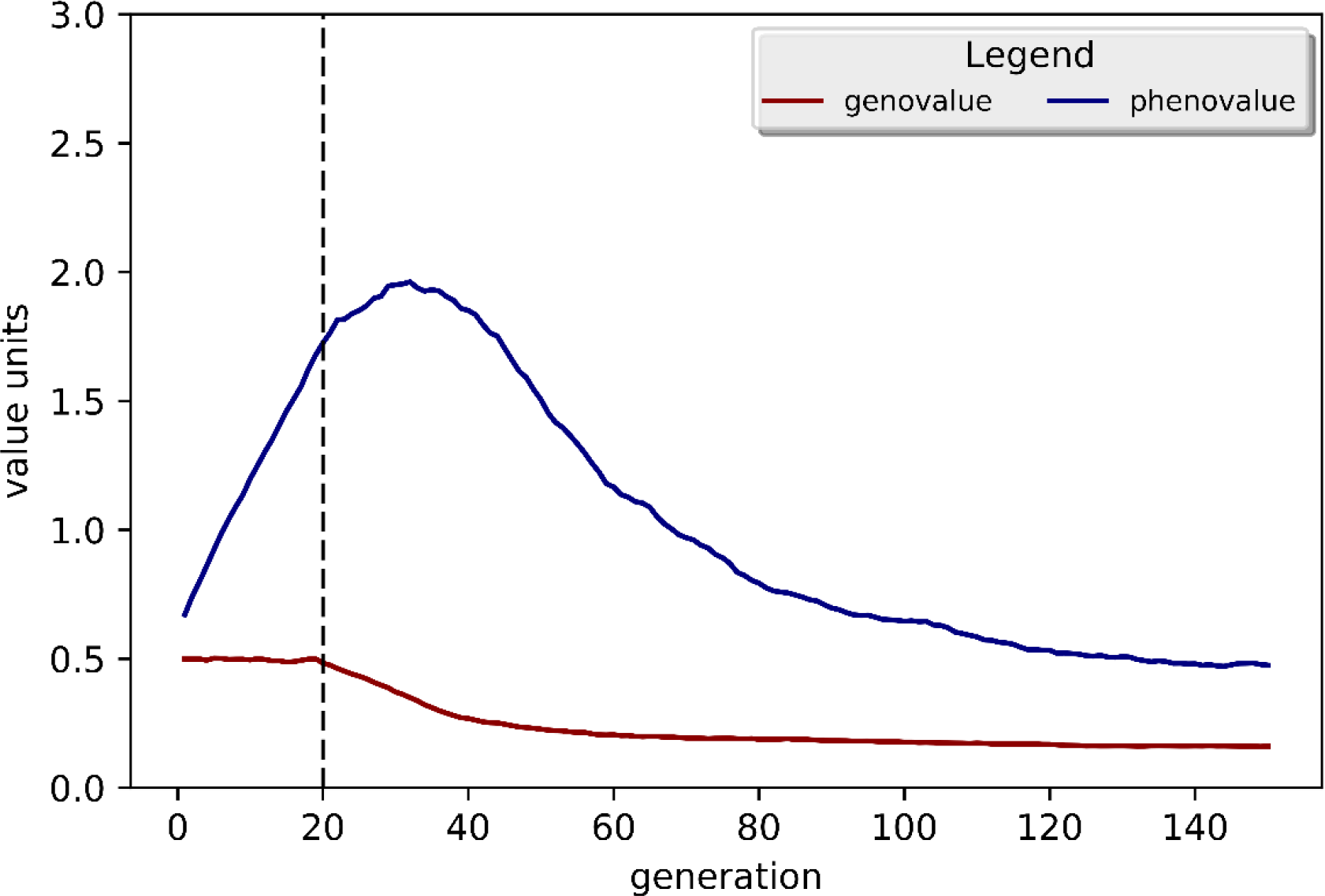
Evolutionary trajectories of EA with a shock at generation 20 (dashed line). The coefficient of effect of phenotype on fitness, γ, changes from 0 to −0.5. The genotypic value remains constant, then starts to decrease immediately at generation 20. The phenotypic value, on the other hand, experiences a lag.

Unlike the “non-shock” condition where the contribution of EA to fitness, γ, is always negative, the first 20 generations for the “shock” condition (γ=0) does not incur genetic evolution as genetic fitness is uncorrelated with phenotype. As expected, we observe no change in genotypic value up to generation 20. However, in the “pre-shock” period, cultural evolution in the form of prestige-biased copying happens which significantly boosts the phenotype of EA. This cultural evolutionary process reduces phenotypic variation in the absence of genetic change, and consequently the diverging pattern only lasts a relatively short period of time (from generation 20 to 30), compared to the non-shock condition (from generation 0 to 20).

### Effect of prestige-biased transmission and other parameters

To systematically examine the effect of other parameters, I thus used the end point (generation 150) phenotypic and genotypic values to represent the quasi long term evolutionary outcomes within this short time window (Figure 3). As expected, most genotypic values are significantly lower at the end of the simulation run than at the beginning due to natural selection. In accordance with the logic of evolution, larger β (more genetic contribution to phenotype) and larger γ (more phenotypic contribution of EA to fitness) increases the strength of natural selection on genes, and when both β and γ are small we barely observe any decline in genotypic value (top left plot, figure 3). Most end-point variation in phenotypic values, on the other hand, is primarily due to the varying degree of prestige-biased transmission. For example, when the ego’s phenotype depends more on the phenotypic value of a cultural model (β = 0.2), the magnitude of prestige-biased transmission φ is positively correlated with the end point phenotypic values. In other words, it “helps” drive up the population mean phenotypic value. Interestingly, an increase in end-point phenotypic value due to larger cultural reproductive skew is accompanied by a smaller end-point genotypic value under certain parameter settings (β = 0.2, β_inter_ = 0.1). This is because the same conditions that leads to large phenotypic variance also leads to large genotypic variance, thus as cultural evolution drives up mean phenotypic value, genetic evolution pushes down mean genotypic value. Expectedly, these conditions are also where we observe the initial diverging patterns, except when φ = 0 (no prestige-based transmission at all).

**Fig 3.**
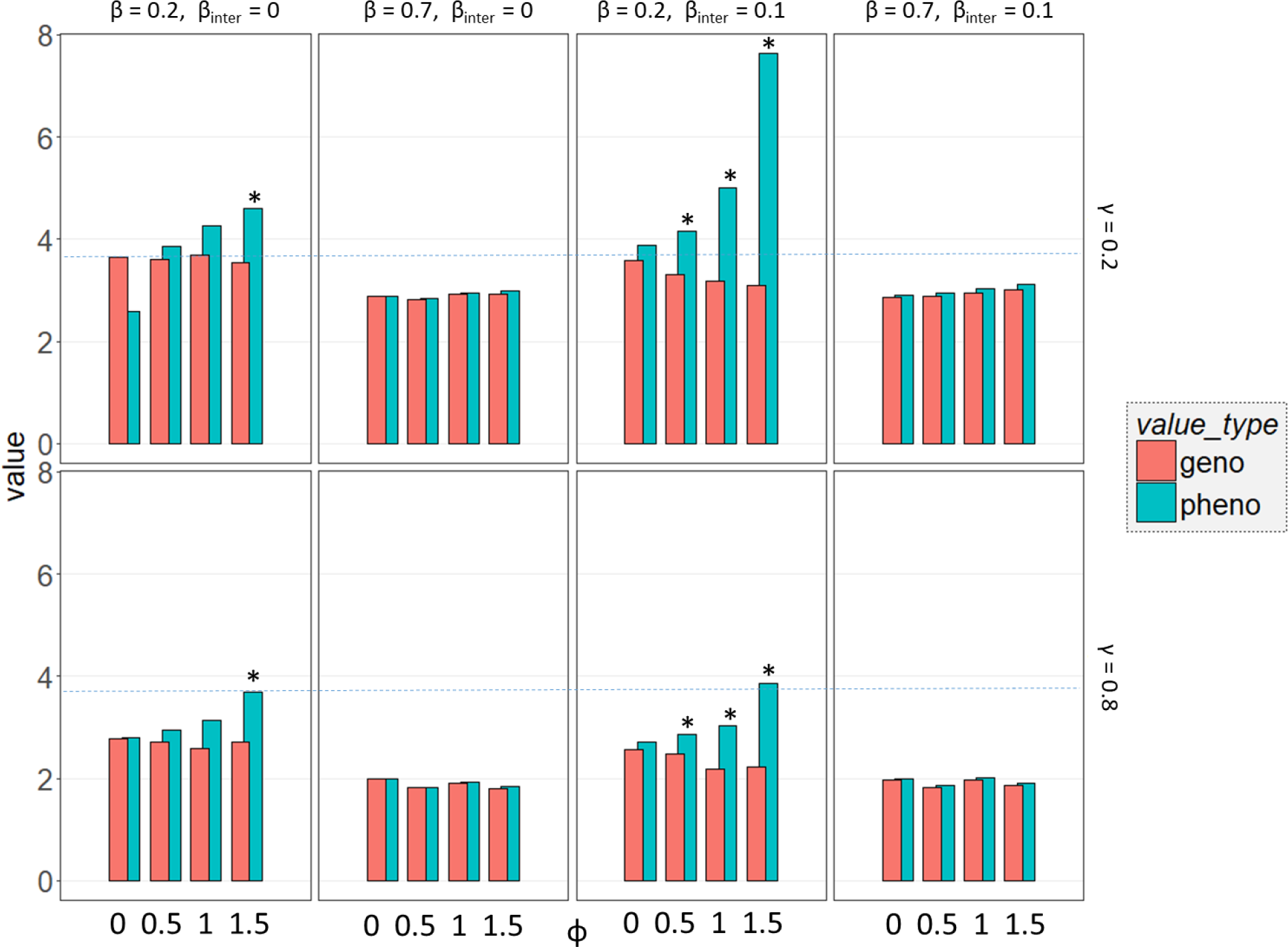
End point phenotypic and genotypic values under different parameter combinations. All data are multiplied by 100 and then log transformed to accommodate for a few extremely large values. Stars represent a definitive diverging pattern of genotypic values and phenotypic values (maximum original phenotypic value >0.9). The dashed lines represent the expected starting genotypic and phenotypic values. Each pair of bars is the average of 1000 independent simulation runs.

## Discussion

### Simulation as a way of studying gene-culture coevolution

It is no news that human cognitive and behavioral traits are significantly influenced by human culture (34), and much effort has been put into modeling the gene-culture coevolutionary dynamics analytically (15, 35). However, most models are designed for analyzing long term allele frequencies and evolutionarily stable scenarios for discrete traits. These models are not ideal for examining short term evolutionary changes, especially when models are becoming more complex from incorporating additional assumptions and parameters.

The evolution of educational attainment serves a case in point. Although it is not difficult to envision a situation where phenotype and genotype undergo different selective pressures, the complexity of real-world biological systems often makes it difficult for humans to evaluate not only the effect of individual assumptions and parameter settings but also the short-term/long term evolutionary outcomes. For example, the steady increase in educational attainment over the past century can be plausibly attributed to income increase (23) or “top-down” government policies aiming at increasing literacy (36), but it can be difficult to know the genetic consequences of these environmental effects or to examine how certain cultural evolutionary mechanisms—such as prestige or payoff biased transmission influence the evolutionary dynamics of EA. In these situations simulation can be of great help. The technique of simulation itself has been widely used by the cultural evolution community to both derive novel insights (37, 38) and provide validation to analytic predictions (39), yet there has been very little simulation attempt to track the evolutionary change of genetic and cultural component of quantitative traits simultaneously. In the present case, the non-intuitive hill-shape phenotypic value change provides important insight for both understanding the role of specific transmission biases in shaping genetic evolution and potentially predicting the short term evolutionary change in the trait of interest (EA).

### Gene and culture evolving towards opposite directions: How should we interpret the result?

There can be many environmental factors that contribute to the phenotypic change of any human trait. In particular, the technological progress and vastly improved living conditions since the industrial revolution has drastically altered many human phenotypes; for example, adult human height has substantially increased throughout the world over the past century due to improved nutrition and reduction in childhood disease (40). What is especially interesting about EA is its consistent negative association with fertility. In general, more educated women delay the onset of childbearing and have fewer children overall compared to less educated women (17, 41). This pattern is very robust in both developed (42) and developing countries (43), and various theories have been proposed to explicate the intrinsic, potentially causal relationship between education level and fertility (44, 45).

This negative association between EA and fertility, which is a reasonable proxy for fitness, has important evolutionary consequences, as both theoretical predictions and empirical evidence suggest that the genetic component of EA is undergoing a temporal decline (22). The phenotype of EA, on the other hand, may experience a very different type of selective pressure. A large literature in sociology shows that educational attainment and socio-economic status are associated (46, 47), and cultural evolutionary theory predicts that humans readily copy the behaviors of the those who are perceived as more prestigious or successful (25, 27). Although educational attainment cannot be “copied” in a literal sense, individuals who pick models with high EA are likely to be more motivated in learning and committed to their academic studies, and as a result become more likely to obtain higher EA themselves. Thus, genetic fitness and cultural fitness of the same trait (EA) invites selection in opposite directions, and what is shown in the simulation model is that biased transmission alone can lead to a short term increase in phenotype.

In reality, the massive increase in EA and over the past century is almost certainly a combination of various social, economic, and cultural factors, and the educational expansion has been described as an “environmental override” (48). The substantial increase in the phenotype of EA indicates that whatever the mechanism is, the decline of genetic component of EA has been vastly overcompensated by enhanced environmental inputs. However, the simulation results reveal a crucial insight: short term change in phenotypic value does not guarantee long-term phenotypic change in the same direction. A natural question to ask is whether there is some ceiling effect of environmental input: that is, whether the environmental factors can drive up the phenotype of EA indefinitely. Unfortunately, the exact relationship between any behavioral genotypes and phenotypes are poorly understood (49). Yet it seems reasonable to assume some upper limit of environmental influences, which means at some point genetics will become the limiting factor in determining the phenotype of EA, which may eventually decline as a result of natural selection.

It has been suggested that because humans constantly modify their environment, the association between education and fertility may not remain negative long enough to have noticeable effects (49). This is certainly a possibility, but at this point we simply do not have enough information to accurately predict future cultural trends. Though it is not difficult to imagine a situation where highly educated, successful people also have more offspring, there is a genuine trade-off between the time one invests in education and in raising offspring. Alternatively, education could be dissociated from status or prestige, yet this possibility does not seem very probable given the great emphasis on education in modern economies (50, 51). A disappearance or reversal in the selection on EA is likely to involve large scale, top-down intervention.

Even in the short term, the decline of the genetic component of EA may have non-trivial consequences. Although people can reasonably question whether educational attainment is the trait under selection (52), it is not crucially important what the precise trait is. What matters in this evolutionary process is that genetic variants associated with EA are being selected against. We know that these genetic variants are primarily involved in brain development processes and neuron-to-neuron communication (14), which accords with previous research showing that the same set of SNPs is associated with both educational attainment and cognitive function (20). Thus, it seems that at least part of the genetic component of EA comes in the form of cognitive ability, which is important for not only EA but also other aspects of social outcomes such as income (53) and job performance (54).

### Are contemporary humans fitness maximizers?

A long-standing question in evolutionary biology is whether organisms behave as if they are maximizing fitness (55). Humans, as an evolved primate, must be subject to the logic of evolution via natural selection (6); yet on the other hand, due to the presence of a second, cultural inheritance system (15), human behavioral evolution seems to require a different type of analysis. Richerson and Boyd (2005) offer a plausible account of how cultural transmission could result in maladaptions –when genetic fitness is not perfectly aligned with cultural fitness for any particular trait. Stated broadly, our capacity for culture evolved so that we could acquire information socially, which conferred significant fitness advantage in the evolutionary past; yet the very same psychological mechanism allows for maladaptations at the same time. In the modern environment, copying the most prestigious individuals may not be the most adaptive strategy (for an extreme case, see De Leo and Heller 2008 on the social transmission of suicide).

However, this view does not go unchallenged by evolutionary biologists. El Mouden et al. (2014) present a narrative of why genetic fitness should always align with cultural fitness by arguing that “genetic selection will shape psychological mechanisms to avoid cultural traits that bear a genetic fitness cost”. This line of thinking assumes an infinite strategy space, in particular the possible psychological mechanisms that can detect genetic fitness consequences in novel environments. Even if we ignore all constraints on the potential evolution of human cognition, it remains the case that novel mutations reaching evolutionary equilibrium typically require some significant amount of time (59).

What is very notable about contemporary humans is that the rate of environmental change can be very rapid. For example. substantially reduced infant mortality (60), dissociation of sexual activity from reproduction (61), and the reversed association between status and fertility (62) happened only in the past few hundred years, and it is unrealistic for genetic evolution to generate new psychological mechanisms to keep up with the cultural change. Thus, to understand the short-term evolutionary dynamics it would be better to focus on what is known about the selective pressure and the heritability of the target of selection in contemporary societies.

## Conclusion

This paper demonstrates the potential utility of a type of simulation model that specifies both the phenotype function and the fitness function in tracking short term evolutionary change in human behavior traits. The results indicate that payoff/prestige biased transmission can significantly affect the evolutionary trajectory of educational attainment in contemporary societies, and that a short term increase due to increase in environmental input towards phenotype may not last indefinitely. The long-term evolutionary trajectory of EA depends on a few key parameters, and researchers can gain valuable insight from analyzing the conditions under which the genotypic value and phenotypic value change as a result of both genetic and cultural selection.

## Acknowledgements

I thank Dr. Joseph Henrich for his support, encouragement, and advice for this project, and Graham Noblit and Nick Patterson for helpful comments and suggestions. This work was supported by Harvard Graduate Training Grant.

